# The Gfr uptake system provides a context-dependent fitness advantage to *Salmonella* Typhimurium SL1344 during the initial gut colonization phase

**DOI:** 10.1101/2025.05.06.652348

**Authors:** Lea Fuchs, Cora Lisbeth Dieterich, Elena Melgarejo Ros, Philipp Keller, Leanid Laganenka, Christopher Schubert, Julia A. Vorholt, Jörn Piel, Wolf-Dietrich Hardt, Bidong D. Nguyen

## Abstract

*Salmonella enterica* serovar Typhimurium (*S*. Tm) is a major cause of foodborne diarrhea. However, in healthy individuals, the microbiota typically restricts the growth of incoming pathogens, a protective mechanism termed colonization resistance (CR). To circumvent CR, *Salmonella* strains can utilize private nutrients that remain untapped by the resident microbiota. However, the metabolic pathways and environmental niches promoting pathogen growth are still not completely understood. Here, we investigate the significance of the *gfr* operon in gut colonization of *S*. Tm, which is essential for the utilization of fructoselysine (FL) and glucoselysine (GL). These Amadori compounds are present in heated foods with high protein and carbohydrate contents, particularly in Western-type diets. We detected FL in both mouse chow and the intestinal tract of mice and showed that *gfr* mutants are attenuated during the initial phase of colonization in the murine model. Experiments in gnotobiotic mice and competition experiments with *Escherichia coli* suggest that *gfr*-dependent fitness advantage is context-dependent. We conclude that dietary Amadori products like FL can support *S*. Tm gut colonization, depending on the metabolic capacities of the microbiota.

## Introduction

*Salmonella enterica* serovar Typhimurium (*S*. Tm), a gram-negative enteric pathogen, is a common cause of foodborne diarrhea (Kirk et al., 2015; Pires et al., 2015). Most *Salmonella*-infected people develop self-limiting gastroenteritis with mild symptoms or remain asymptomatic whereas immunocompromised persons, young children and the elderly are at risk of developing a severe systemic infection (Gal-Mor et al., 2014; Pang et al., 1995). The factors determining why some individuals remain asymptomatic while others develop diarrheal disease are not yet fully understood. Mouse models have provided valuable insights into the molecular mechanisms of *S*. Tm infections and colonization resistance (CR), a microbiota-driven defense against pathogens (Herzog et al., 2023; Stecher et al., 2013; Tsolis et al., 2011). In healthy animals with a complex microbiota, CR typically restricts pathogen growth in the gut lumen, preventing *S*. Tm colonization in most cases - only 5% of the infected mice exhibit detectable colonization (Kreuzer & Hardt, 2020; Stecher & Hardt, 2011). Contrarily, mouse models with defined but a less complex microbiota have reduced CR, leading to consistent colonization. These gnotobiotic mouse models include Oligo-MM^12^ and low complexity microbiota (LCM) mice, whose microbiota consists of 12 or 8 strains, respectively (Brugiroux et al., 2016; Hoces et al., 2023; Stecher et al., 2010). *Salmonella* spp. overcome CR by utilizing a large variety of metabolic pathways enabling growth on at least 100 different carbon sources, some of which may be unused by the host’s resident microbiota (Gutnick et al., 1969; Seif et al., 2018; Warsi et al., 2019). We hypothesize that such private nutrients may promote *S*. Tm colonization, consistent with previous studies showing that galactitol or arabinose utilization can enhance *S*. Tm luminal growth (Eberl et al., 2021; Gul et al., 2023; Ruddle et al., 2023). Previously a random-barcoded transposon sequencing (RB-TnSeq) screen identified *S*. Tm genes *gfrA* (SL1344_4465) and *gfrB* (SL1344_4471) as important for colonizing the LCM mice (Nguyen et al., 2020; Nguyen et al., 2024). These genes are part of the *gfrABCDEF* operon, which encodes a mannose family phosphotransferase system (PTS) important for fructoselysine (FL) and glucoselysine (GL) import and degradation (Miller et al., 2015) (**Fig. 1A**). FL/GL utilization systems can be found in other species, e.g. *Enterococcus faecium* and *E. coli* (Graf von Armansperg et al., 2021; Miller et al., 2015; Wiame et al., 2002). Specifically, *gfrABCD* encodes a PTS permease that transports and phosphorylates both FL and GL while *gfrE* and *gfrF* encode for deglycases that cleave glucoselysine 6-phosphate and fructoselysine 6-phosphate, respectively (Miller et al., 2015) (**Fig. 1A**).

**Figure 1.**
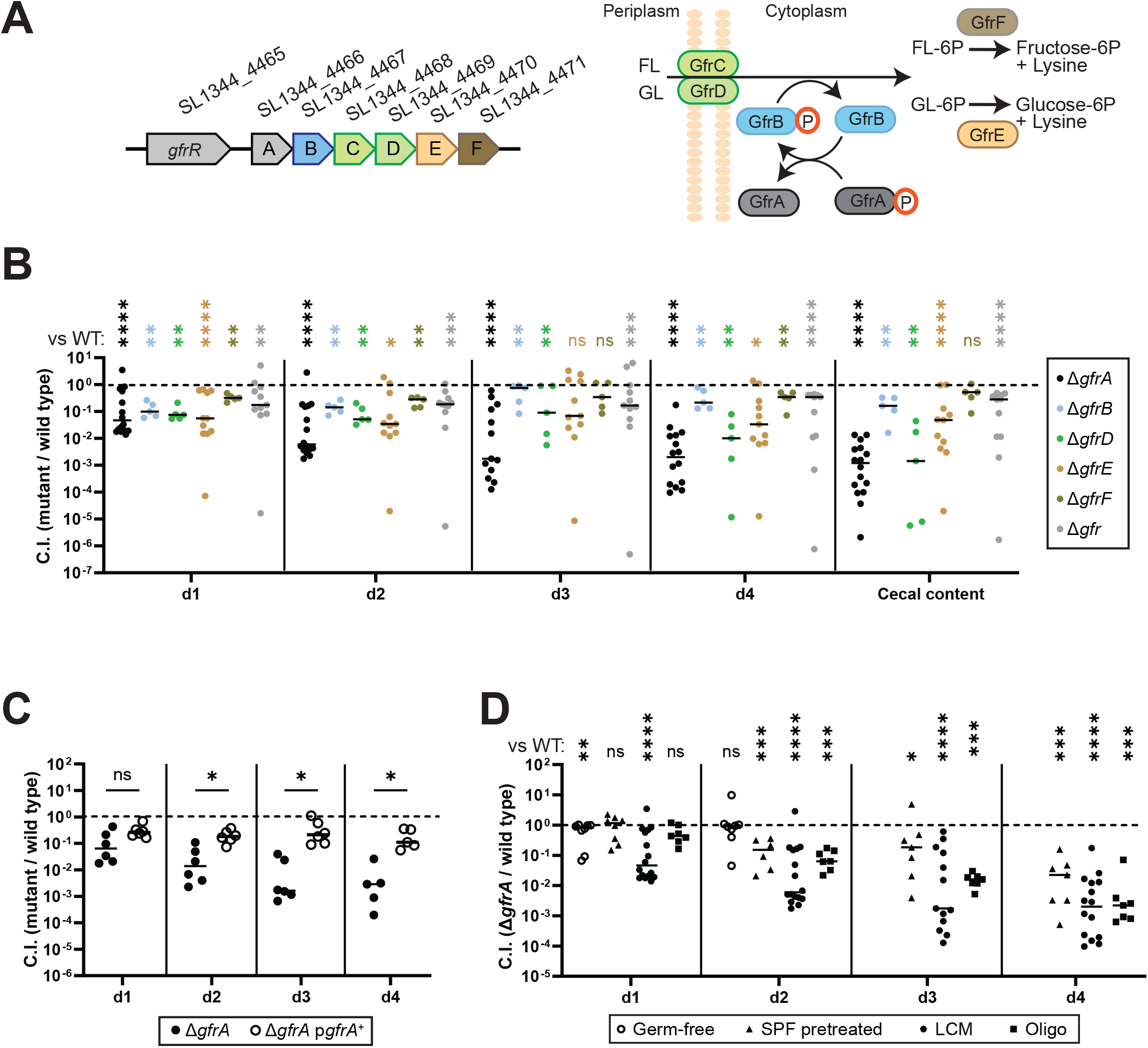
The fructoselysine transport system supports gut luminal *S*. Tm growth during initial growth in OligoMM^12^ and LCM mice. (**A**) *gfr* operon and proposed function of the Gfr enzymes. GfrC and GfrD function as EIIC and EIID PTS subunits, forming the transporter to facilitate FL and GL import across the membrane, while GfrA and GfrB act as EIIA and EIIB subunits, respectively, mediating phosphoryl transfer. The deglycases GfrE and GfrF cleave fructoselysine-6P or glucoselysine-6P, respectively, yielding glucose-6P or fructose-6P and lysine. Modified after (Miller et al., 2015). (**B-D**) Mice were orally infected with an equal mixture of *S*.Tm SL1344 strains at a titer of 5 × 10^6^ CFU in total. Each mixture contained a reference wild-type strain. (**B**) Competitive infections of *gfr* subunit deletion strains (Δ*gfrA*, Δ*gfrB*, Δ*gfrD*, and Δ*gfrF*), *gfr* operon deletion (Δ*gfr*), and wild type *S*. Tm in LCM mice. qPCR analysis of chromosomal WITS tags was used to determine the competitive index (C.I.). **(C)** Competitive infection comparing WT, Δ*gfrA*, and a complemented Δ*gfrA* strain (carrying a plasmid-encoded copy of *gfrA*) in LCM mice. Strain abundance was analyzed by differential plating of fecal and cecal content on MacConkey agar. (**D**) Competitive infection of Δ*gfrA* and wild-type strain in germ-free, streptomycin-pretreated SPF, LCM, and OligoMM^12^ mice. (**B-D**) Each point represents one mouse, at least 5 mice per group. The Mann-Whitney U test was used for statistical analysis; *p < 0.05; **p < 0.01; *** p < 0.001; **** p < 0.0001.

Amadori compounds are formed by the Maillard reaction during heating or dehydration of fruits and vegetables (Ames, 1992; Baynes et al., 1989; Erbersdobler, 1977). Most Amadori compounds, including FL and GL, are not consumed by the host and can only be metabolized by a limited number of gut microbiota strains (Erbersdobler et al., 1970). Thus, FL or GL might represent private nutrients that could support the growth of *S*. Tm strains. Recent studies have investigated the impact of FL on the gut microbiota. FL found in formula milk was shown to enrich for FL degradation genes in multiple taxa in formula-fed infants (Erbersdobler & Faist, 2001; Martysiak-Żurowska & Stołyhwo, 2007; van Dongen et al., 2022). In adults, individual variations in microbial FL degradation have been reported (van Dongen et al., 2021), suggesting that different microbiotas feature different capacities to utilize FL. We hypothesized that the LCM mouse microbiota lack such FL consumers, which could explain why *gfrA* and *gfrB* mutants were attenuated at gut luminal colonization (Nguyen et al., 2020; Nguyen et al., 2024). *S*. Tm 14028 has been shown to utilize FL and, to a lesser extent, GL as carbon or nitrogen source in *in vitro* experiments (Miller et al., 2015) and to rely on fructose-asparagine, another Amadori product, for growth in the inflamed gut (Ali et al., 2014). However, based on differences in other carbohydrate utilization genes (i.e. its distinct ability to use galactitol (Eberl et al., 2021; Gul et al., 2023; Ruddle et al., 2023)), it remained unclear if *S*. Tm SL1344 would similarly rely on FL or GL utilization. Here, we investigated whether FL serves as a private nutrient for *S*. Tm SL1344 during colonization of various murine models.

## Results

### The *gfr* genes of *S*. Tm are required for initial growth in the gut lumen of mice

Attenuated gut-luminal growth of *S*. Tm *gfrA* and *gfrB* mutants was observed in a genetic screen of *S*. Tm SL1344 mutants in LCM mice (Nguyen et al., 2020; Nguyen et al., 2024). Notably, LCM mice exhibit two distinct infection stages: an initial phase during days 1–2 post-infection (p.i.), characterized by pathogen growth in the presence of an undisturbed microbiota, followed by an inflammation-associated bloom (Nguyen et al., 2020; Nguyen et al., 2024). To verify the colonization defect of *gfrA* and *gfrB* mutants and to assess the *in vivo* contributions of the additional genes within the *gfr* operon, we constructed single gene in-frame deletion mutants for *gfrA, gfrB, gfrD, gfrE* and *gfrF* and a deletion mutant lacking the entire *gfrABCDEF* operon (Δ*gfr*) in *S*. Tm SL1344 (**Figure 1A**; **Table 1**). Each strain was labeled with a unique strain-specific DNA barcode sequence (WITS tag), enabling strain fitness measurements via qPCR (Grant et al., 2008; Nguyen et al., 2020).

LCM mice were co-infected with an equal mixture of wild type, Δ*gfrA*, Δ*gfrB*, Δ*gfrD*, Δ*gfrE*, Δ*gfrF* and Δ*gfr* strains (in total 5x10^6^ CFU, by oral gavage; **Fig. 1B**). Fecal pellets were collected on days 1, 2, 3 and 4 post infection (p.i.) to quantify the competitive fitness of each mutant. The Δ*gfrA* mutant exhibited the strongest attenuation compared to the wild-type strain with a competitive index (C.I.) of 0.1-0.01 by day 1 p.i., declining further to 0.001 by day 4 p.i. (**Fig. 1B**). The Δ*gfrB*, Δ*gfrD*, Δ*gfrE*, Δ*gfrF* and Δ*gfr* mutant strains were attenuated by about 3 to 10-fold (C.I. ≈ 0.1) on day 1 p.i. and either remained at this level of attenuation (Δ*gfrB*, Δ*gfrE*, Δ*gfrF*, Δ*gfr*), or declined further to C.I. of 0.1-0.01 (Δ*gfrD*) by day 4 p.i.. These data suggest that the *gfr* operon provides *S. Tm* with a competitive advantage during initial growth in LCM mice, as the Δ*gfr* and the individual *gfr* subunit mutants except for the Δ*gfrA* mutant were primarily attenuated during the first two days of infection.

The additional attenuation observed in the Δ*gfrA* mutant was not due to second-site mutations, as the phenotype could be complemented by introducing a wild-type copy of *gfrA* under its native promoter on a plasmid. In competition with a wild-type strain, the complemented mutant grew to near wild-type levels in LCM mice (**Fig. 1C**), confirming that the colonization defect is specifically attributable to the *gfrA* deletion. We speculate that the previously described regulatory function of EIIA enzymes in other PTS systems (Maze et al., 2014) might explain why the Δ*gfrA* mutant exhibits more pronounced attenuation than the other subunit deletion strains or the whole operon deletion strain.

In an effort to provide a foundation for future work on *gfrA*, we have assessed the context dependence of *gfrA*’s role in mice harboring different microbiotas (**Fig. 1D)**. Specific pathogen-free (SPF) mice pretreated with antibiotics have lower colonization resistance compared to gnotobiotic LCM and Oligo-MM^12^ mice, due to abundant free monosaccharides and a significantly reduced gut microbiota (Barthel et al., 2003; Garner et al., 2009; Ng et al., 2013; Nguyen et al., 2024; Schubert et al., 2025; Stecher et al., 2005). Infection kinetics also differ; in streptomycin-pretreated mice, inflammation occurs as early as day 1 p.i., without a distinct initial growth phase, in contrast to the delayed inflammatory response observed in LCM or Oligo-MM12 mice (Kreuzer & Hardt, 2020; Nguyen et al., 2024). In streptomycin pretreated SPF mice, the attenuation of Δ*gfrA* strain in a competitive infection is less pronounced than in LCM or Oligo-MM^12^ mice (Brugiroux et al., 2016; Kreuzer & Hardt, 2020) (**Fig. 1D**). In germ-free (GF) mice with even higher monosaccharide availability (Schubert et al., 2025), the Δ*gfrA* mutant showed no significant fitness loss for at least 2 days (C.I.≈1; **Fig. 1D**). Since GF mice are particularly sensitive to *S*. Tm infection (Stecher et al., 2005), we have limited that experiment to 2 days p.i.. These data indicate that the microbiota composition affects the degree of attenuation of the Δ*gfrA* mutant. They also suggest that FL or GL is particularly important if other nutrients are scarce. Taken together, the *gfr* operon confers a fitness advantage in a context dependent manner.

### Fructoselysine is present in mouse chow and the mouse intestine

Amadori products are primarily found in food and are not known to be synthesized by the host or gut commensals. Since Amadori products cannot be absorbed by the host (Tuohy et al., 2006), we anticipated that FL and GL would accumulate in the gut lumen in sufficient quantities to support *S*. Tm growth. Given that GL has been described as a poor nutrient source for *S*. Tm 14028, we focused our analysis on FL (Miller et al., 2015). Additionally, the chemical synthesis of GL has been reported to produce substantial amounts of Heyns side products, which would complicate the isolation of pure GL for analysis. Mass spectrometry performed on extracts from mouse food pellets and mouse intestinal content from LCM mice verified the presence of FL in both the diet and the intestine. The total ion chromatograms (TIC) of food pellet and LCM mouse intestine extracts revealed a complex mixture of compounds (**Fig. 2**). The extracted ion chromatogram (EIC) specific to FL confirmed its presence in both the food pellet extract and the contents of the gut lumen of uninfected LCM mice.

**Figure 2.**
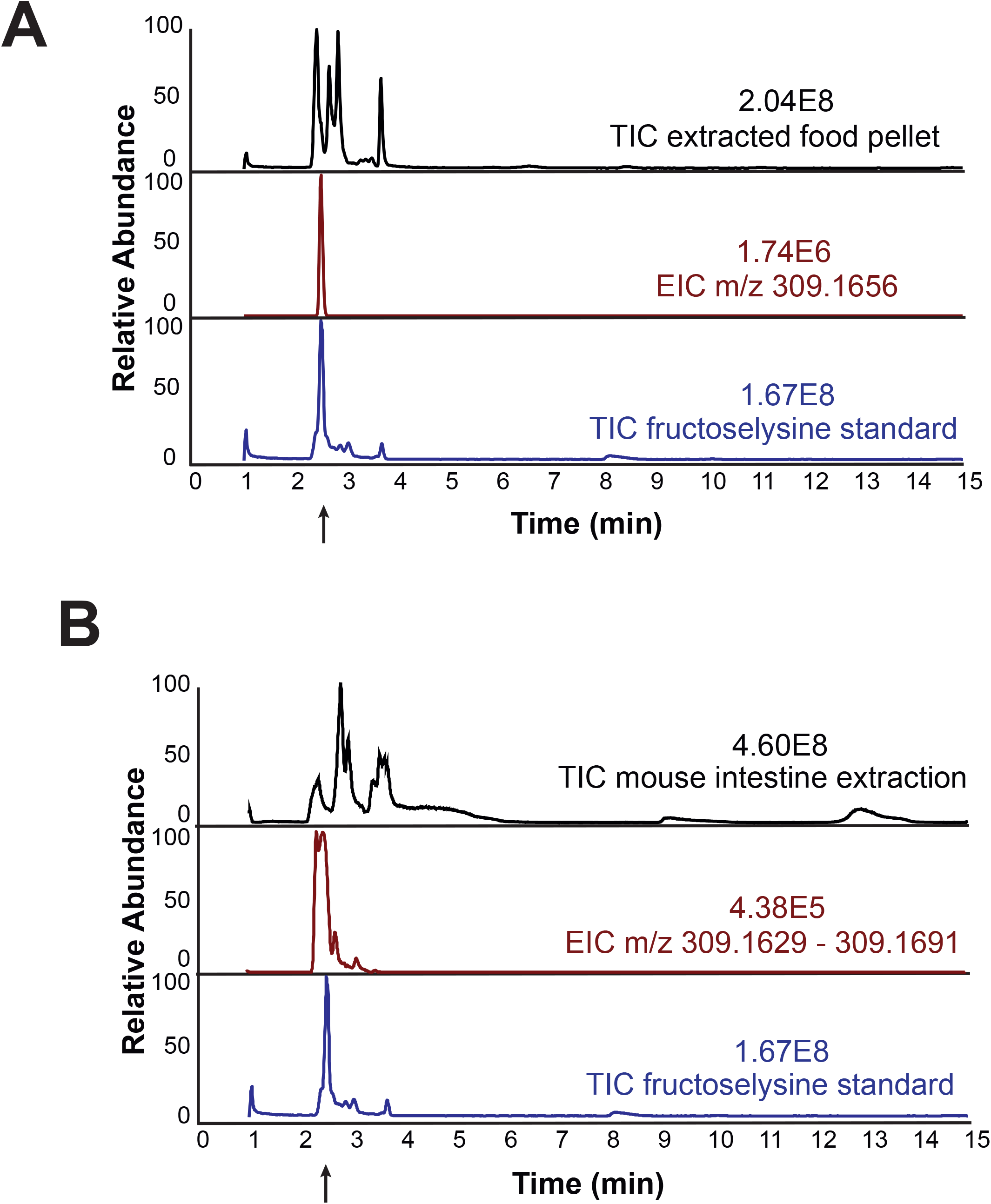
Fructoselysine was detected in food pellets and in the intestine of LCM mice. (**A**) Upper panel: total ion chromatogram (TIC) of whole food pellet extraction in water; middle panel: extracted ion chromatogram (EIC) of FL specific m/z 309.1668; lower panel: TIC of FL standard. (**B**) Upper panel: TIC of whole mouse intestine extraction in water, middle panel: EIC of FL specific m/z 309.1629 – 309.1691, lower panel: TIC of FL standard.

### Fructoselysine feeding does not enhance growth of *Salmonella* in the gut

Next, we investigated whether FL supplementation could enhance *S*. Tm growth in the mouse gut or compromise colonization resistance in SPF mice with an intact conventional microbiota. To test this, SPF mice were pretreated with a single dose of FL (66 mg or 120 mg) or PBS as a negative control by gavage, followed by oral *S*. Tm infection 4 h later (**Fig. 3A**). The time interval between FL administration and infection was chosen to expose the pathogen cells to elevated FL levels in the gut lumen based on previous studies (Wotzka et al., 2019). No significant differences were observed between the control group and mice administered 66 mg or 120 mg FL (**Fig. 3A**), indicating that the introduction of excess FL to the gut lumen via gavage does not impact *S*. Tm colonization in SPF mice.

**Figure 3.**
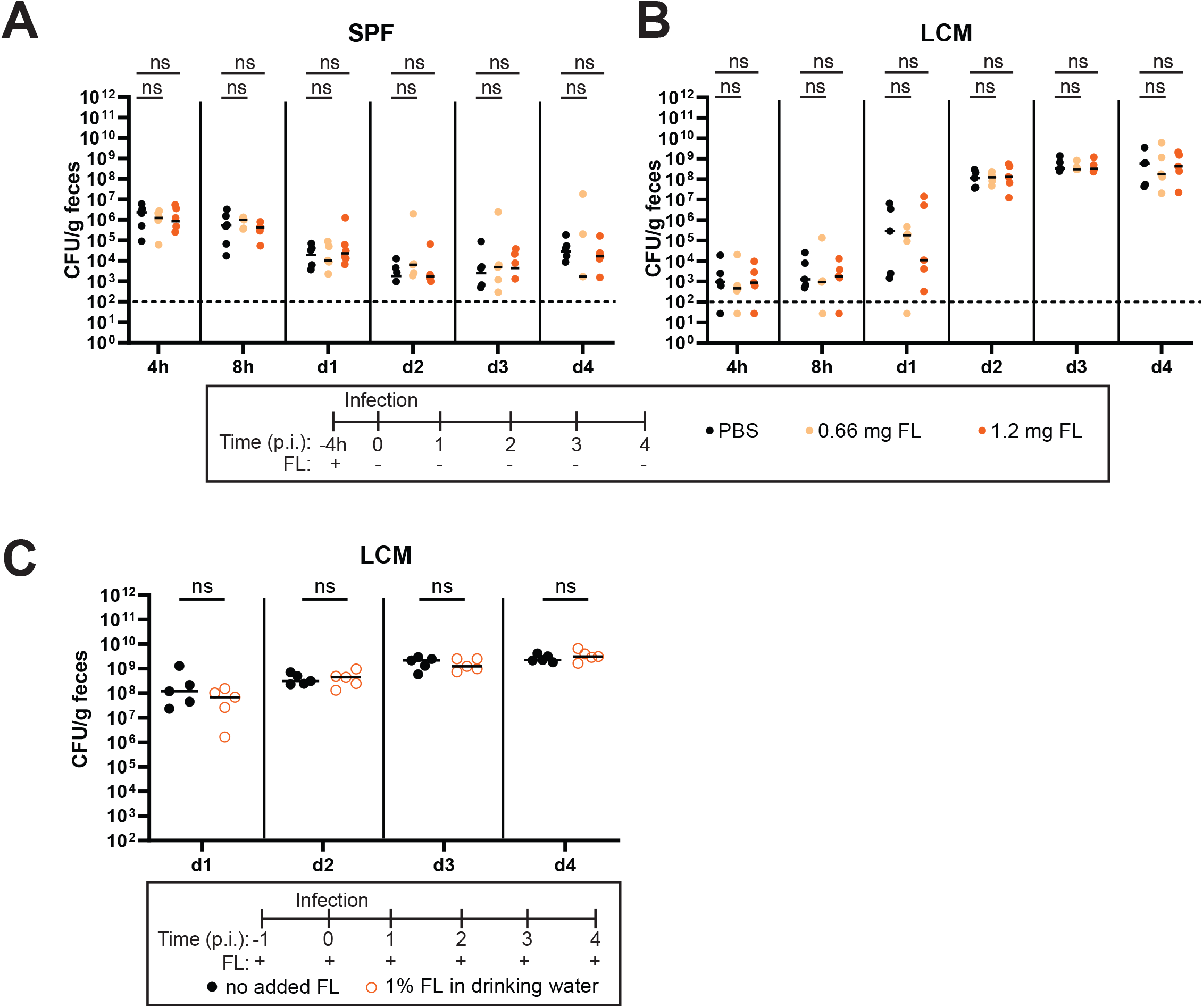
FL supplementation neither suppresses colonization resistance in SPF mice nor enhances luminal growth of *S*. Tm in LCM mice. Fecal loads from 4 h to 4 days p.i. of *S*. Tm in (**A**) SPF or (**B**) LCM mice gavaged with 66 mg or 120 mg of FL or control (phosphate buffered saline, PBS) 4 h before infection. (**C**) Fecal loads of *S*. Tm in LCM mice provided with 1% FL in drinking water compared to control mice with normal water. Pathogen loads were enumerated by plating dilutions of feces on MacConkey agar. Each point represents one mouse (N = 5 per group from two independent experiments each).

We reasoned that SPF mice lack key nutrients, making FL supplementation alone insufficient to support *S*.Tm growth. Since *S*.Tm can colonize LCM mice without antibiotic perturbation, we further reasoned that such limiting nutrients are more readily available in LCM than in SPF mice, allowing FL supplementation to further enhance growth. However, LCM mice receiving a single dose of 66 mg or 120 mg or a continuous supply of 1% FL in drinking water showed similar *S*. Tm loads in feces compared to the control groups (**Fig. 3B and C**). We conclude that FL is already sufficiently abundant in our mouse food and that other factors are limiting further gut-luminal growth of *S*. Tm, at least under the conditions tested in our mouse experiments.

### Fructoselysine is a carbon and nitrogen source for *S*. Typhimurium SL1344 under anaerobic conditions

FL can serve as a carbon and nitrogen source in culture under aerobic conditions for *E. coli* and *S*. Tm 14028 (Miller et al., 2015; Wiame et al., 2002), but for SL1344, this remained unknown. Given the importance of *gfr* genes for gut luminal growth, we aimed to assess the capacity of SL1344 to utilize FL under anaerobic conditions with fumarate, a key electron acceptor in the murine gut (Nguyen et al., 2020). We also aimed to verify the role of *gfrA* in FL utilization, as well as the substrate-specific functions of *gfrE* (GL cleavage) and *gfrF* (FL cleavage). The wild type and mutant strains exhibited no significant growth differences in minimal medium (M9 with fumarate) supplied with glucose and ammonium as sole carbon and nitrogen sources, respectively, reaching a maximum OD_600_ of ≈ 0.6 (**Fig. 4A**). This suggests that there are no inherent general growth defects in the mutant strains. However, when the carbon source was switched to FL, both the wild type (WT) and Δ*gfrE* strains exhibited significantly reduced growth compared to glucose, with both reaching a final OD_600_ of ≈ 0.15. The Δ*gfrA* knockout strain exhibited no growth, confirming the critical role of *gfrA* in FL utilization (**Fig. 4B**). The Δ*gfrF* mutant was similarly expected to show no growth. However, its residual growth in media with FL as the sole carbon source may suggest that GfrE has a relaxed specificity, allowing it to cleave FL at a reduced rate or that an alternative deglycase can cleave fructoselysine-6-phosphate. All strains exhibited similar growth patterns in minimal medium with FL as the sole nitrogen source (and pyruvate as additional carbon source) (**Fig. 4B**), though growth was lower, which suggests that FL is a less effective source of nitrogen. Overall, our *in vitro* growth experiments indicate that FL is a relatively poor nutrient source when provided as the sole carbon or nitrogen source, especially in comparison to glucose or ammonium.

**Figure 4.**
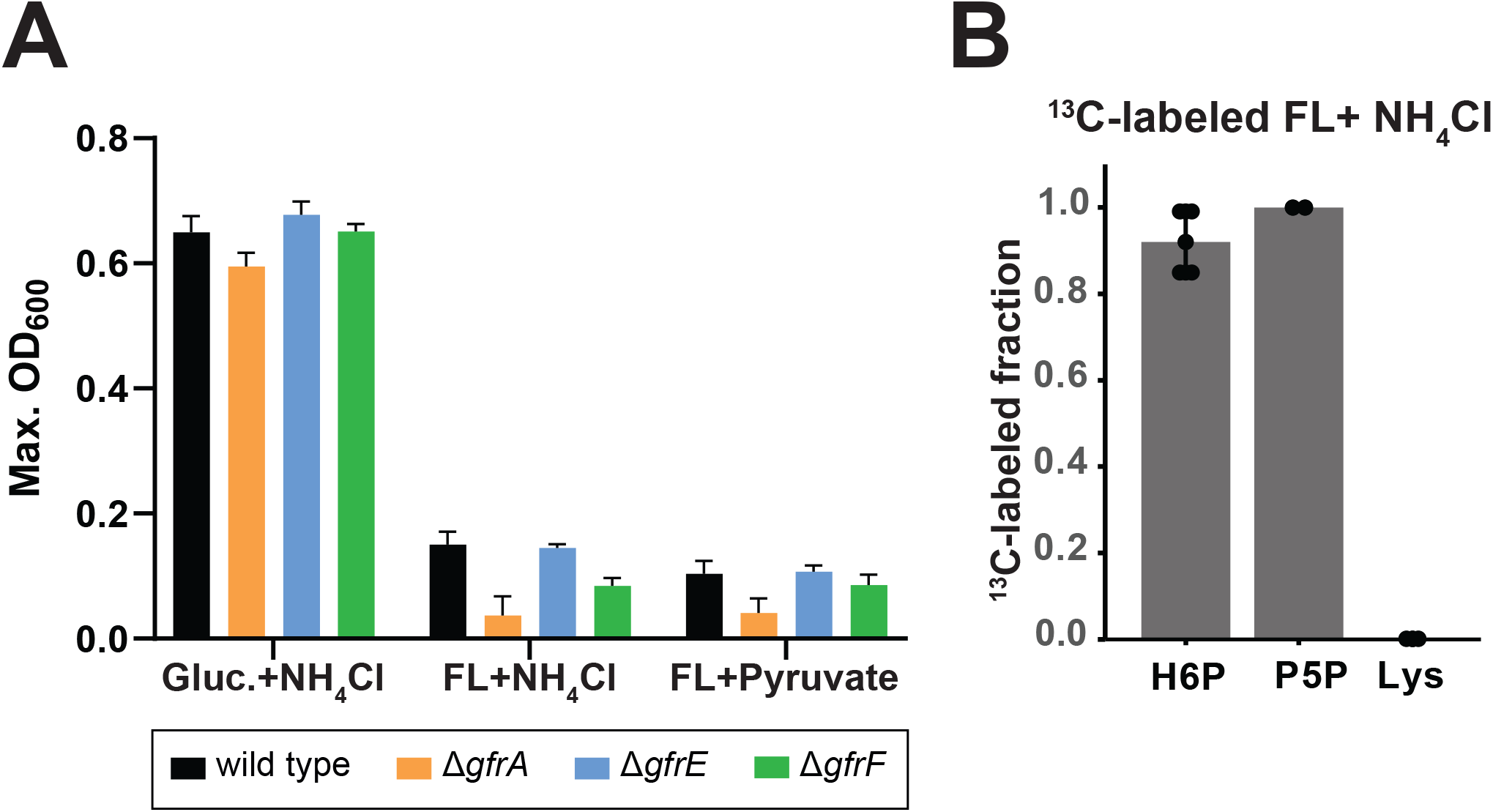
FL fails to support robust *in vitro* growth as the sole carbon or nitrogen source. **(A)** Anaerobic growth of *S*. Tm WT, Δ*gfrA*, Δ*gfrE and* Δ*gfrF* in minimal medium (M9 with 312.42 µM fumarate) supplemented with one of the following: (left) 22 mM glucose and 19 mM NH_4_Cl, (middle) 22 mM FL as a sole carbon source with 19 mM NH_4_Cl, or (right) 18 mM FL as a sole nitrogen source with 44 mM pyruvate. Data represent mean ± SEM from two independent experiments (N = 6). The maximum optical density (OD) was recorded over the course of 800 minutes. (**B**) ^13^C-labeled fraction of the indicated metabolites in wild type *S*. Tm grown in M9 medium supplemented with 22 mM FL with a fully labeled fructose moiety and an unlabeled lysine moiety and 19 mM NH_4_Cl (N = 3; H6P= Hexose 6-phosphate; P5P= Pentose 5-phosphate; Lys= lysine).

To determine how *S. Tm* utilizes carbon from both the carbohydrate and lysine moieties of FL, we performed ^13^C-tracing experiments using FL with a fully ^13^C-labeled fructose-moiety and unlabeled lysine-moiety. To ensure a metabolic steady state, cells were grown anaerobically for at least five generations in minimal M9 medium containing the partially labeled FL and ammonium before sampling. During exponential growth, intracellular metabolites were extracted, and the labeled fractions of hexoses-phosphates, pentose-phosphates, and lysine were quantified by LC-MS. Hexose- and pentose-phosphate molecules were fully labeled, indicating that they are derived from the ^13^C-labeled fructose moiety via glycolysis and the non-oxidative branch of the pentose phosphate pathway, respectively (**Fig. 4C**). Endogenous lysine remained unlabeled, indicating that it originated from the unlabeled lysine moiety of FL. This suggests that carbons from the fructose moiety of FL were not used for *de novo* lysine synthesis and that lysine-moiety can be directly utilized as a substrate for protein synthesis. Overall, these results indicate that the fructose moiety of FL, can be catabolized although FL supported significantly slower growth compared to the preferred carbon source glucose.

### Fructoselysine provides lysine as a building block for protein biosynthesis

Previous work has identified ammonium and asparagine as preferred nitrogen sources for *E. coli* (Schubert et al., 2021). This preference also holds true for *S*. Tm SL1344, as our results demonstrate that FL is a poor nitrogen source compared to ammonium and asparagine for anaerobic growth on minimal media (**Fig. 4A and Fig. 5A**). These data suggest that FL may not be effective for nitrogen assimilation. To determine whether this contributes to FL being a poor nitrogen source, we assessed lysine’s capacity to serve as a nitrogen source. Interestingly, L-lysine supported even less growth than FL (**Fig. 5A; Supplementary Fig. 1**). To determine whether FL can rescue lysine auxotrophy, we analyzed the growth of a lysine auxotrophic mutant strain (Δ*lysA*) in medium supplemented with FL or L-lysine. This mutant strain lacks the ability to synthesize L-lysine, making import from external sources essential for growth. Under aerobic conditions, FL supplementation rescues the growth of the Δ*lysA* mutant in minimal media supplied with glucose and ammonium in a dose dependent manner (**Fig. 5B**). While L-lysine promoted auxotrophic growth more effectively than FL at lower concentrations (0.1-1.0 mM), both nutrients supported comparable growth at higher concentrations (≥ 5 mM) (**Fig. 5B**). These observations suggest that the limited capacity of FL to support growth under nitrogen-depleted conditions is due to the high energetic cost of lysine degradation and subsequent deamination. Moreover, the ability of FL to rescue lysine auxotrophy is less effective than supplementing the growth medium directly with L-lysine.

**Figure 5.**
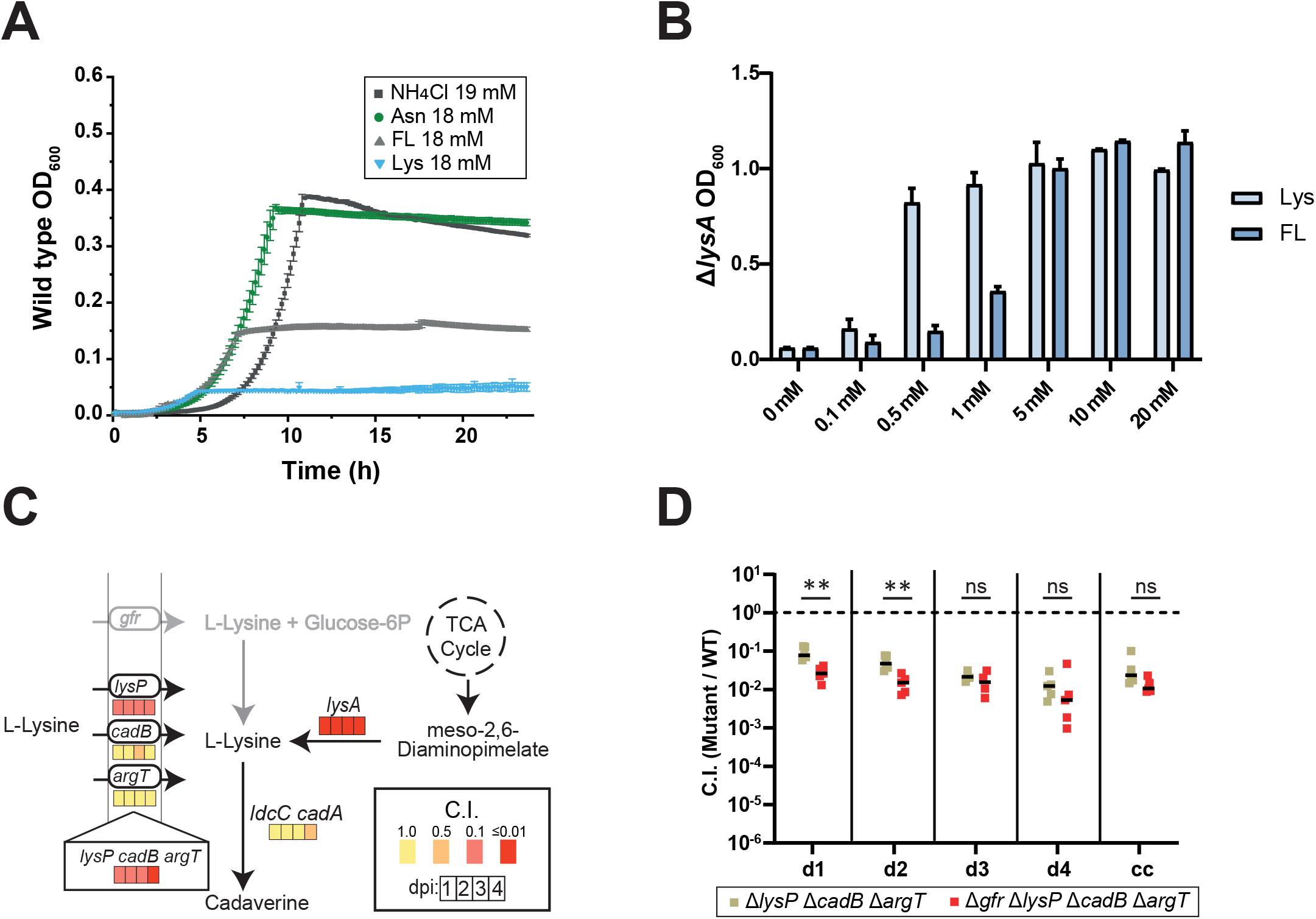
FL can restore growth of lysine auxotrophic strains. (**A**) Wild type *S*. Tm growth under anaerobic conditions in minimal medium (M9 with 312.42 µM fumarate) supplemented with 22 mM glucose and various nitrogen sources (NH_4_Cl, asparagine (Asn), fructoselysine (FL), and lysine (Lys); concentrations indicated in legend) (N = 3). (**B**) Growth of the lysine auxotrophic strain (Δ*lysA*) in minimal medium supplemented with 22 mM glucose, 19 mM NH_4_Cl, and various concentrations (0-20mM) of either lysine or FL under aerobic conditions. Mean OD_600_ values were recorded at 12 hours of growth (N = 5, from two independent experiments). (**C**) Mean competitive indexes (C.I.) of DNA-barcoded *S*. Tm mutant strains deficient in lysine transporters (LysP, CadB, ArgT), lysine biosynthesis (LysA), and lysine catabolic enzymes (LdcC and AadA), relative to WT strains (median fitness) in fecal samples from five Oligo-MM12 mice collected from day 1 to day 4 p.i.. Colored boxes indicate the degree of attenuation for day 1-4 p.i.. (**D**) Competitive indices of lysine transport mutants (ΔlysP, ΔcadB, ΔargT) compared to an isogenic strain with *gfr* operon deletion (Δ*gfr*. Δ*lysP* Δ*cadB* Δ*argT)*. Abbreviations: cc, cecal content on day 4 p.i.. Statistical analysis was performed using the Mann-Whitney U test; *p < 0.05, **p < 0.01.

Together, the *in vitro* growth and *in vivo* competition assays suggest that FL provides a source of lysine, which may be limiting in the gut. This is supported by the significant attenuation of Δ*lysP* Δ*cadB* Δ*argT* triple mutant, which is defective in lysine uptake, in OligoMM^12^ mice (C.I.: 0.1–0.01; **Fig. 5C**), indicating *S*.Tm’s dependency on exogenous lysine. Our data also reveal that LysP is the primary lysine importer, as the Δ*lysP* mutant was strongly attenuated (C.I.: 0.1), whereas Δ*argT* and Δ*cadB* mutants grew similarly to the wild type. However, lysine importers ArgT and CadB contribute modestly during late-stage infection, as deficiency of both further attenuates the Δ*lysP* mutant on day 4 post-infection (**Fig. 5C**). Exogenous lysine is insufficient, as lysine biosynthesis appears critical for growth (**Fig. 5C**; C.I. of Δ*lysA* mutant: 0.01). These findings indicate that lysine is limiting, and that FL may serve as an additional lysine source for gut luminal growth of *S*. Tm. Indeed, Δ*gfr* deletion further attenuates the Δ*lysP* Δ*cadB* Δ*argT* triple mutant (compared Δ*lysP* Δ*cadB* Δ*argT vs* Δ*gfr* Δ*lysP* Δ*cadB* Δ*argT* in **Fig. 5D**) during initial growth, suggesting that GL/FL contribute to the cellular L-lysine pool required for growth.

As lysine is an ineffective nitrogen donor, we expect that its degradation would not be critical for growth. Consistent with this, the Δ*ldcC* Δ*cadA* mutant, which is incapable of lysine decarboxylation, the first step in lysine degradation, exhibited wild type levels of growth in the murine gut (**Fig. 5C**).

### *S*. Tm and *E. coli* compete for fructoselysine in the gut

The detection of FL in the gut content of LCM mice suggests that LCM microbiota lack FL consumers. This raises the question of whether a direct competitor for FL or GL could influence *S*. Tm colonization in LCM mice. Some *E. coli* strains can utilize FL via the *frl* operon (Laganenka et al., 2023; Wiame et al., 2002). Although the *frl* system belongs to a different class of transporters, we hypothesized that it may enable *E. coli* to compete with *S*. Tm SL1344 in an FL-dependent manner. To investigate a potential competition between *E. coli* and *S*. Tm for FL *in vivo*, LCM mice were pre-colonized with either *E. coli* WT Z1331, a human isolate, or its FL-utilization deficient mutant strain (Δ*frl*) 24 h prior to *S*. Tm infection. On day 1 and day 2 p.i., wild-type *E. coli* suppressed *S. Tm* gut colonization more efficiently than *E. coli* Δ*frl* strain (**Fig. 6A**). However, this difference disappeared at later time points, further supporting that the FL utilization plays a critical role only during the initial growth phase of the infection. *S. Tm* growth suppression was not due to a fitness defect of the Δ*frl* strain, as both *E. coli* strains reached comparable colonization levels in the gut of LCM mice (**Fig. 6B**).

**Figure 6.**
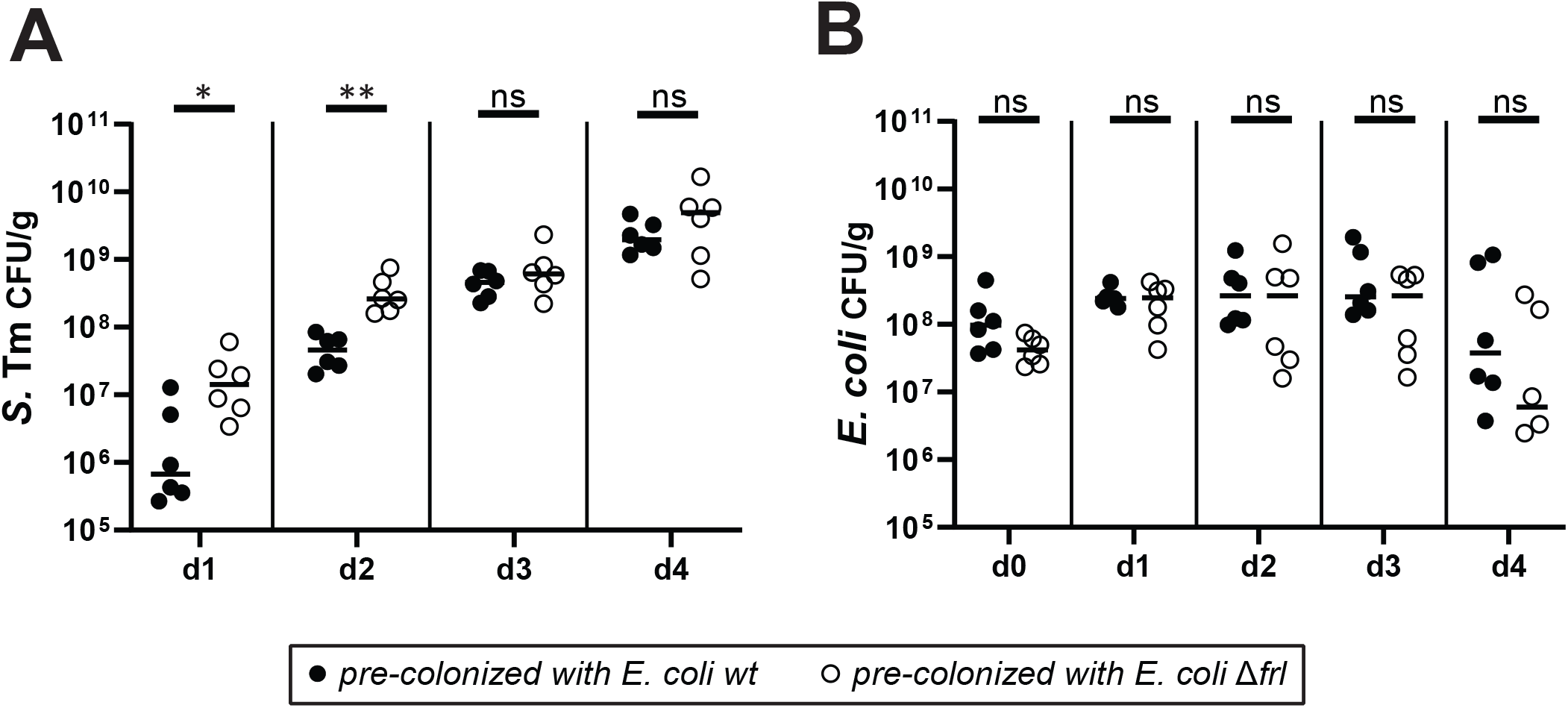
*E.coli* suppresses *S*. Tm during initial growth in a FL-utilization dependent manner. Fecal loads of (**A**) *S*.Tm and (**B**) *E. coli* in infected LCM mice pre-colonized with wild type *E*.coli or congenic *frl* deficient *E.coli* strain (Δ*frl*). Mice were orally gavaged with 5x10^7^ CFU *E. coli* one day before receiving 5x10^7^ CFU *S*.Tm by oral gavage. Fecal loads were determined by plating fecal samples on McConkey agar. Each point represents one mouse. The Mann-Whitney U test was used for statistical analysis; *p < 0.05; **p < 0.01.

## Discussion

Understanding the interplay between dietary compounds and gut colonization by enteric pathogens is crucial, as nutrient availability directly influences microbial competition and pathogen growth. While the effects of Amadori products on the microbiota and pathogens have gained attention in recent years, their role in promoting pathogen colonization is not fully understood (Ali et al., 2014; Rampanelli et al., 2025; Seiquer et al., 2014). In this study, we showed that the Amadori product FL is important for gut colonization by *S*. Tm strain SL1344, particularly during the initial growth phase. Data from the various mouse models suggest *gfr*-mediated gut luminal growth is context-dependent, notably under conditions of low nutrient availability. Unlike fructose-asparagine, which plays a major role during later stages of infection when inflammation is pronounced (Ali et al., 2014), we find that *gfr* mutants are attenuated during the initial growth, when *S*. Tm competes with an intact microbiota (**Fig. 1B**). The *gfr* operon is less critical in GF mice or in streptomycin pre-treated animals, where other nutrients are more abundant (Nguyen et al., 2024; Schubert et al., 2025) (**Fig. 1D**). This context-dependent fitness benefit is reminiscent of growth advantages *S*. Tm SL1344 or certain *E. coli* strains gain from galactitol utilization (Eberl et al., 2021; Gul et al., 2023; Ruddle et al., 2023). Both galactitol and FL utilization require redox balancing, which limits their use under anaerobic conditions, explaining why galactitol and FL utilization become relevant only in specific competitive contexts. Overall, our findings reinforce the idea that *S*. Tm growth is shaped by nutrient availability, microbiota composition and the host’s food choices.

Our data indicate that FL is not an optimal nutrient source, as it fails to support robust growth when supplied as the sole source of either carbon or nitrogen. While the carbohydrate moiety of FL can be catabolized anaerobically, the lysine moiety likely contributes only to protein biosynthesis, as its degradation has multiple oxidation steps (Knorr et al., 2018) that are constrained by the reducing environment of the gut. Thus, it is not a surprise that neither FL nor lysine serves as an efficient sole nitrogen source in our anaerobic growth assays (**Fig. 4A and Fig. 5A**). These limitations likely explain why additional FL supplementation did not promote *S*. Tm blooms in SPF and LCM mice. However, FL-derived lysine can serve as a source for lysine *in vitro* (**Fig. 5B**). Lysine appears to be a limiting nutrient in the gut, as both import and biosynthesis are required for growth (**Fig. 5C**). Thus, *in vivo*, FL serves as a general carbon source and, in addition, contributes to the cellular lysine pool for protein biosynthesis. In a complementary approach, we could show that the presence of diet-derived FL in the gut of LCM mice supported *S*. Tm growth in a *gfr*-dependent manner, due to the absence of competing FL/GL consumers. It has been reported that FL can be consumed by some *Enterobacteriaceae* in the mammalian gut (Erbersdobler et al., 1970; Erbersdobler & Somoza, 2007; Wiame et al., 2002). Our mouse experiments with *E. coli* revealed that FL uptake systems contribute to competition between different enterobacteriaceal strains *in vivo*. Specifically, we showed that *E. coli* can suppress *S*. Tm in an FL-utilization dependent manner during early colonization (**Fig. 6**). Recently, FL was identified as a chemoattractant involved in niche segregation in *E. coli*, suggesting that FL sensing may influence competitive behavior. This points to a complex crosstalk between the *E. coli* Frl system, fructose-lysine metabolism, and autoinducer-dependent quorum sensing (Laganenka et al., 2023).

In conclusion, we showed that fructose-lysine uptake via the Gfr system plays a context-dependent role for *S*. Tm during initial growth in the colonized gut. The presence of a direct competitor can reduce colonization, suggesting that diversifying the gut microbiota, along with dietary alterations, may help to fortify colonization resistance against invading pathogens. Investigating the influence of dietary components on pathogen colonization can provide valuable insights on strengthening natural gut defenses and the developing new strategies to prevent *S*. Tm infections.

## Materials and Methods

### Bacterial strains, growth conditions, and reagents

The strains and plasmids used in this study are listed in **Supplementary Table 1**. The *E. coli* strains are derived from *E. coli* Z1331, a motile stool isolate from healthy human volunteer (Wotzka et al., 2018). All strains were routinely grown overnight at 37°C in Lysogeny broth (LB) supplemented with ampicillin (100 µg ml^−1^), kanamycin (50 µg ml^−1^), streptomycin (50 µg ml^−1^) or chloramphenicol (15 µg ml^−1^) with agitation. For the *in vitro* growth assays, M9 minimal medium (47.75 mM Na_2_HPO_4_·7H_2_O, 22.04 mM KH_2_PO_4_, 8.55 mM NaCl, 100 µM CaCl_2_, 2 mM MgSO_4_) supplemented with trace elements (13.4 mM EDTA, 3.1 mM FeCl_3_-6H_2_O, 0.62 mM ZnCl_2_,76 nM CuCl_2_-2H_2_O, 42 nM CoCl_2_-2H_2_O, 162 nM H_3_BO_3_, 8.1 nM MnCl_2_-4H_2_O) and 312.42 µM fumarate was used as a base medium. Bacteria were grown overnight in the base medium with 22 mM glucose and 19 mM NH4Cl under aerobic (ambient atmosphere) or anaerobic conditions (gas atmosphere 7% H_2_, 10% CO2, 83% N_2_). Samples were washed twice in ddH_2_O to remove residual medium and resuspended in fresh base medium supplemented with the indicated nitrogen and carbon sources. Bacterial suspensions were adjusted to OD_600_= 0.02. The OD was monitored for 24 hours at 37°C under constant orbital shaking using a Biotek Synergy H1 (SH1M2-SN) under aerobic or anaerobic conditions as indicated. For the histidine auxotrophic strain *S*. Tm SL1344, media were supplemented with 1.6 mM of L-histidine.

### Construction of mutant strains

The Δ*gfrA*, Δ*gfrB*, Δ*gfrE* and Δ*gfrF* mutants were constructed by P22 phage transduction from a *S*. Tm 14028 mutant library (Porwollik et al., 2014) into a SL1344 strain background (Hoiseth & Stocker, 1981). The Δ*gfr* and Δ*gfrD* mutant strains were constructed using the lamda red recombination system (Datsenko & Wanner, 2000). The whole *gfr* operon and *gfrD* gene were either replaced with the *cat* cassette from pKD3 or the *aphT* cassette from pKD4 by homologous recombination (oligonucleotides used for this purpose are listed in **Supplementary Table 1**). Mutated alleles were subsequently transduced into a fresh strain background by P22 phage transduction. DNA barcodes from wild-type isogenic tagged strains (WITS) or wild-type isogenic standardized hybrid (WISH) strains were used to individually tag all wild-type and mutant strains via P22 phage transduction, followed by selection on their respective antibiotics (Daniel et al., 2024; Grant et al., 2008). All constructs were verified by PCR and whole genome sequencing. All strains and their corresponding genotype are listed in **Supplementary Table 1**.

### Animals

All mice were reared at the EPIC facility at ETH Zürich. C57BL/6J mice (Jackson Laboratory, catalogue no. JAX:00066) were held under specific pathogen-free conditions in individually vented cages (light:dark cycle 12:12 h, room temperature 21 ± 1 °C, humidity 50 ± 10%). LCM mice are ex-germ-free C57BL/6J mice that have been associated with the strains of the altered Schaedler flora (Maier et al., 2013; Stecher et al., 2010). Oligo-MM^12^ mice are ex-germ-free C57BL/6J mice, which were colonized with a defined consortium of 12 bacterial strains isolated from the murine gut (Brugiroux et al., 2016). All gnotobiotic and germ-free (GF) mice were bred in flexible film isolators under strict exclusion of microbial contamination. The 8–12-week-old mice of both sexes were randomly assigned to the experimental groups. All animal experiments were reviewed and approved by Tierversuchskommission, Kantonales Veterinäramt Zürich under license ZH158/2019 and ZH109/2022, complying with the cantonal and Swiss legislation.

### Fructoselysine synthesis

Fructoselysine was synthesized as described previously (Miller et al., 2015). Labeled fructoselysine was synthesized from D-glucose-^13^C_6_.

### Mass spectrometry analysis of fructoselysine in food pellets and mouse intestine

Cecal content from euthanized LCM mice was collected in a non-autoclaved collection tube, immediately flash-frozen and lyophilized afterwards. 5 mg of lyophilized cecal content or food pellet was dissolved in 50 µl MS grade water. After vortexing vigorously, the samples were incubated at room temperature for 15 minutes and separated by centrifugation for 10 minutes at maximal speed. 45 µl of the supernatant was transferred into autosampler vials with a built-in 100 µl insert. MS spectrometry measurements were performed on a Dionex Ultimate 3000 HPLC system coupled to a Thermo ScientificTM Q ExactiveTM Hybrid Quadrupole-Orbitrap Mass Spectrometer. Experiments were conducted using solvents A (H2O + 0.1 % formic acid) and B (MeCN + 0.1 % formic acid) with a Phenomenex Kinetex 2.6 µm XB-C18 150 x 4.6 mm. The used method contained a flowrate 1 mL/min, with the gradient of 1% B 0-4 min, 1% to 4% B 4-10 min, 4% to 95% B 10-13 min, 95% B 13-14 min and 95% to 1% B 14-15 min. MS-settings: Spray Voltage 3.5 kV, capillary temperature 320 °C, sheath gas (52.50), aux gas (13.75), spare gas (2.75), probe heater 437.50 °C, S-Lens RF (50), positive mode, resolution 70.000, AGC target 3e6, microscans 1, maximum IT 200 ms, scan range 50-750 m/z.

### Metabolic conversion of 13C-labeled fructoselysine

#### Growth conditions

*S*. Tm cultures were grown in triplicates under anaerobic conditions for at least five generations in M9 minimal media containing 22 mM labeled fructoselysine (^13^C fructose, ^12^C lysine) as carbon source and 19 mM NH_4_Cl as nitrogen source. *S*. Tm was first grown for 12h in M9 media with glucose and NH_4_Cl under anaerobic conditions. Subsequently, it was subcultured into M9 medium supplemented with labeled fructoselysine and grown until mid-exponential phase, followed by a second passage in the same labeled medium. At mid-exponential growth phase, 0.4 OD units of culture (1 mL at OD_600_= 0.4) were applied to a 0.2 µm polyethylene filter (prewashed with 50 °C warm ultra-pure water (MilliQ)) under vacuum and then washed with 4 mL 37 °C warm ultra-pure water (MilliQ). After washing, filters were transferred into 8 mL precooled quenching solution (60: 20: 20, acetonitrile: methanol: 0.1 M formic acid, −20 °C.) and incubated for 15 min on ice to extract intracellular metabolites. Extracts were snap-frozen in liquid-nitrogen and lyophilized at −40 °C. Dried samples were reconstituted in ice-cold ultra-pure water (MilliQ) to a biomass concentration of 400 ng/µL (assuming that 1 OD unit has 250 µg cell dry weight) and further diluted in acetonitrile to a final biomass concentration of 100 ng/µL.

#### Liquid-chromatography mass-spectrometry

Metabolites were analyzed using ultra-high pressure liquid chromatography (UPLC Ultimate 3000, ThermoFisher Scientific, Reinach, Switzerland) equipped with a hydrophilic interaction liquid chromatography (HILIC) column (Agilent Infinity Lab Poroshell 120 HILIC-Z; 100 Å; 2.7 µm; 100 x 2.1 mm; Peek-lined SS, Agilent, Basel, Switzerland) coupled to a hybrid quadrupole-orbitrap mass spectrometer (Q Exactive Plus, ThermoFisher Scientific, Reinach, Switzerland). The solvent system as described in Hsiao et al., 2018 but without medronic acid consisted of 10 mM ammonium acetate in MilliQ (solvent A) and 10 mM ammonium acetate in 90% acetonitrile (solvent B). For metabolite separation the following gradient was used at a constant flow rate of 500 μL / min: 10% A for 1 min; linear increase to 40% A over 5 min; 40% A for 3 min; linear decrease to 10% A over 0.5 min; and equilibration at 10% A for 3.5 min. Samples were measured in negative and positive mode. Mass spectrometry settings were: Fourier transform mass spectrometry with a spray voltage of –2.8 kV (negative); 2.8 kV (positive), a capillary temperature of 275 °C, S-lens RF level of 50, an auxiliary gas flow rate of 20, and an auxiliary gas heater temperature of 350 °C. Mass spectra were recorded as centroids at a resolution of 70’000 at a mass to charge ratio (m/z) of 435 with a mass range of 75–800 m/z and a scan rate of ≈4 Hz in full scan mode. 10 µL of sample corresponding to 1 µg of biomass were injected. LC-MS measurements were analyzed by emZed3 (Kiefer et al., 2013). Metabolites were identified by m/z (mass tolerance of 0.004 mass units) and by retention time. Retention time was confirmed by commercially available standards. The peak area cut off was set at 5000/µg biomass. Isotopologue fractions (s_i_) and labeled fraction (LF) were calculated as described before (Buescher et al., 2015) by targeted peak integration of all detected isotopologues according to Equation 1 and Equation 1. Calculated values were corrected for natural ^13^C labeling.

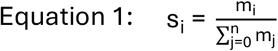

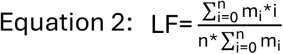

### Mouse infections and competitive infection experiments

The 8-to-12-week-old SPF mice were orally pretreated with streptomycin (50 mg) 24h before infection. No pretreatment was required before the infection of GF, LCM and Oligo-MM^12^ mice. *S*. Tm and *E. coli* cultures were grown 12 h in LB containing the appropriate antibiotics at 37°C with shaking, diluted 1:20 and sub-cultured for 4h in the same medium without antibiotics. Bacteria were washed twice in PBS (137 mM NaCl, 2.7 mM KCl, 10 mM Na2HPO4 and 1.8 mM KH2 PO4) and mice were infected with bacteria by gavage (5*10^7^ cfu in 50 µl for SPF mice, 5×10^6^ cfu in 50 µl for LCM and Oligo-MM^12^ and 5*10^4^ cfu for GF). For competitive index (C.I.) experiments mice were infected with 1:1 mixtures of the indicated strains. Fecal samples homogenized in 500 µl PBS at the indicated timepoints were plated on MacConkey agar with streptomycin to quantify fecal *S*.Tm loads. At day 4 post-infection (p.i.), mice were euthanized by cervical dislocation, and cecal contents, liver and spleen segments, and mesenteric lymph nodes were collected and homogenized in 500 µl of PBS with 0.1% Tergitol using a TissueLyser (Qiagen) prior to plating on MacConkey agar to determine *S*. Tm loads. For competitive experiments, barcoded strains were enriched by incubating 250 µl of the homogenate in 3 ml LB containing 50 µg/ml streptomycin for 4 h. Genomic DNA from enrichment cultures was extracted using the QIAamp® DNA Mini Kit (QIAGEN). Relative densities of the different WITS-tagged strains were determined by real-time PCR with tag-specific primers, while WISH-tagged strains were quantified by Illumina sequencing (see below). Competitive indices were normalized to the relative abundance of each strain in the input inoculum. In the competition experiments with *E. coli*, mice were pre-colonized with 5×10^7^ cfu in 50 µl PBS of wild type or the knockout *E. coli* strains 24 h prior to *S*. Tm infection.

#### Illumina Sequencing

WISH-barcoded *S*. Tm experiments were followed by amplicon sequencing to quantify strain abundance, with sample preparation conducted as previously described (Schubert et al., 2025). Illumina sequencing was performed at the BMKGENE (Germany).

### Fructoselysine feeding experiments

SPF or LCM mice were pretreated 4 h before infection with 100 μl of fructoselysine solution in low (0.66 g/ml) or high (1.2 g/ml) concentration. Control mice received PBS. Fructoselysine solutions were prepared in tap water and sterile filtered using a 0.22 μm filter. For FL supplementation in drinking water, 2.5 g of fructoselysine was added to 250 ml of tap water and sterilized by filtration through a 0.22 µm filter. This 1% concentration was based on previous studies on similar sugars, such as galactitol (Sousa et al., 2017) . Mice received fructoselysine-supplemented water starting one day prior to infection and maintained throughout the course of the experiment.

## Supporting information

Supplementary Table 1 Strains_plasmids_primers

## Acknowledgements

We would like to thank members of the Hardt, the Vorholt and the Piel lab for helpful discussions. We acknowledge the staff of the ETH Zürich mouse facility EPIC/RCHCI for expert support of our animal experiments (especially Manuela Graf, Katharina Holzinger, Dennis Mollenhauer, Sven Nowok & Dominik Bacovcin) and the staff of the Microbiology Institute for technical support. This work has been supported by grants from the Swiss National Science Foundation (310030_192567) to WDH and the NCCR Microbiomes (51NF40_180575) funded by the Swiss National Science Foundation to WDH and JV. C.S is supported by the German Research Foundation (SCHU 3606/1-1).

