## Supplementary Figure 1 for "The Gfr uptake system provides a context-dependent fitness advantage to *Salmonella* Typhimurium SL1344 during the initial gut colonization phase"

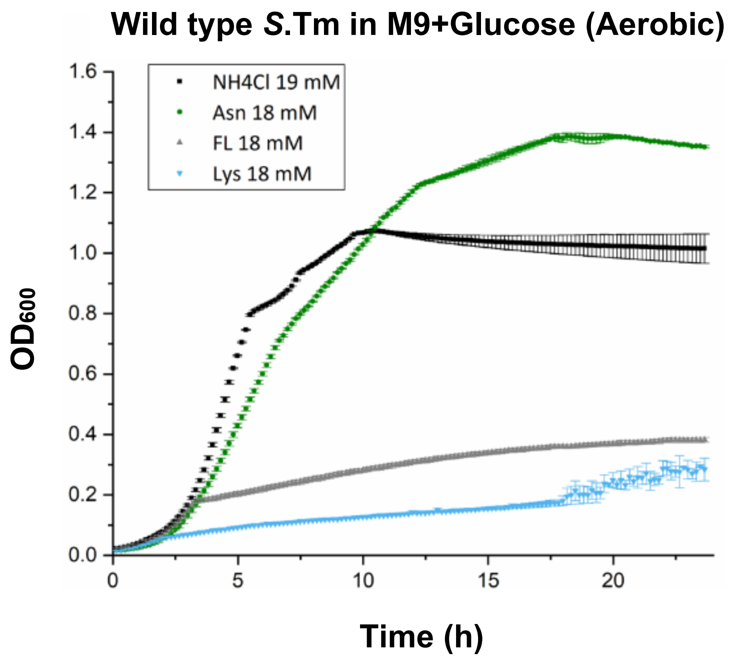


**Supplementary Figure 1:** Wild type *S*. Tm growth under aerobic conditions in M9 medium supplemented with 22 mM glucose with the indicated nitrogen sources (N = 3).
